# SCF^Fbxw5^ targets MCAK in G_2_/M to facilitate ciliogenesis in the following cell cycle

**DOI:** 10.1101/2021.01.15.426831

**Authors:** Jörg Schweiggert, Gregor Habeck, Sandra Hess, Felix Mikus, Klaus Meese, Shoji Hata, Klaus-Peter Knobeloch, Frauke Melchior

**Author notes:** correspondence should be addressed to J.S. or F.M. ( or).

## Abstract

The microtubule depolymerase Kif2C/MCAK plays important roles in various cellular processes and is frequently overexpressed in different cancer types. Despite the importance of its correct abundance, remarkably little is known about how MCAK levels are regulated in cells.

Using comprehensive screening on protein microarrays, we identified 161 candidate substrates of the multi-subunit ubiquitin E3 ligase SCF^Fbxw5^, including MCAK. *In vitro* reconstitution assays demonstrate that MCAK and its closely related orthologs Kif2A and Kif2B become efficiently polyubiquitylated by neddylated SCF^Fbxw5^ and Cdc34, without requiring preceding modifications. In cells, SCF^Fbxw5^ targets MCAK for proteasomal degradation specifically during G_2_/M. While this seems largely dispensable for mitotic progression, loss of Fbxw5 leads to increased MCAK levels at basal bodies, which impair formation of primary cilia in the following G_1_. We have thus identified a novel regulatory event of ciliogenesis that occurs already within the G_2_/M phase of the preceding cell cycle.

## Introduction

In the past decade, primary cilia have gained massive attention due to the discovery of their involvement in a plethora of human diseases, ranging from organ-related disorders such as polycystic kidney disease to more pleiotropic syndromes like Bardet-Biedl^1–3^. By forming antenna-like membrane protrusions that enrich diverse cellular receptors, primary cilia are important signalling hubs for tissue development and homeostasis^4^. Ciliogenesis takes place in almost all human cells and requires drastic remodelling of centrosomes that includes their migration to the cortex and a microtubule-dependent generation of a plasma membrane protrusion^5,6^. Upon re-entry into the cell cycle, primary cilia must be resorbed in order to release centrosomes for spindle formation and thus reappear only after mitotic exit^7^.

Recently, the ubiquitin proteasome system (UPS) has been identified as an important regulator of ciliogenesis^8–10^. Ubiquitylation defines the attachment of ubiquitin to substrate proteins via an enzymatic cascade, in which the final transfer to target lysine residues is catalysed by a so-called E3 ligase in a highly specific manner^11–13^. Since ubiquitin itself contains acceptor lysine residues for other ubiquitin moieties, chains of different linkages can be formed, some of which lead to proteasomal degradation^14^. Examples for ubiquitin-dependent control of ciliogenesis comprise the E3 ligase Mindbomb1 that antagonises ciliogenesis by targeting Talpid3, and the Cullin-RING ligases (CRL) Cul3-KCTD17 and SCF^Fbxw7^ that promote ciliogenesis via degradation of trichoplein and Nde1, respectively^15–17^.

CRLs constitute the biggest family of ubiquitin E3 ligases being responsible for up to 20% of proteolytic ubiquitylation events in cells^18,19^. The best-studied class of CRLs are SCF (Skp1-Cul1-F-box protein) E3 ligases that use Cul1 as a central scaffold protein. Cul1 recruits a small RING (really interesting new gene) containing protein called Rbx1 via its C-terminus and a substrate receptor module composed of Skp1 and one out of 69 interchangeable F-box proteins via its N-terminus^20,21^. Catalytic activity of SCF E3 ligases requires the modification of Cul1 with the ubiquitin-like modifier Nedd8. This induces conformational changes that facilitate ubiquitin transfer from the Rbx1-bound E2-Ubiquitin thioester to the substrate recruited by the F-box protein^22,23^. Fbxw5 belongs to the WD40-domain containing family of F-box proteins. So far, known substrates of SCF^Fbxw5^ include the actin remodeller Eps8, the centrosomal scaffold protein Sas6, the COPII component Sec23b and apoptosis signal-regulating kinase 1 (Ask1)^24–27^. In addition, Fbxw5 has been suggested to be also active within a Cul4A/DDB1-containing CRL, in which it targets the tumour suppressors DLC1 and Tsc2^28,29^.

In order to discover further substrates of Fbxw5, we now employed comprehensive substrate screening on protein microarrays and identified Kif2C/MCAK (mitotic centromere-associated kinesin) as an important novel target of SCF^Fbxw5^. MCAK is the best studied member of the kinesin-13 family, which are non-motile kinesins that use their central motor domain to depolymerise microtubules (MTs) in an ATP-dependent manner^30–32^. MCAK is mostly known for its role in mitosis, where it localises to spindle poles, spindle MTs and centromeres exerting crucial roles in spindle formation and chromosome segregation^33–35^. Its activity is heavily regulated through phosphorylation by different mitotic kinases, such as AuroraA, AuroraB, Cdk1 and Plk1^36–40^. Recently, MCAK has been implicated also in non-mitotic events, such as DNA damage repair and ciliogenesis^41,42^.

Here, we show that SCF^Fbxw5^ regulates MCAK protein levels by specifically catalysing K48 ubiquitin chains on MCAK in a highly efficient manner. Although this process occurs during G_2_/M, loss of Fbxw5 does not provoke severe defects in mitosis but instead impairs ciliogenesis later in the following G_0_ phase. Our work thus demonstrates an intriguing regulatory process required for ciliogenesis that takes place not during the event of ciliogenesis itself but rather within the preceding cell cycle.

## Results

### *In vitro* ubiquitylation screening identifies 161 candidate substrates of SCF^Fbxw5^

In order to identify novel substrates of the SCF^Fbxw5^ complex, we performed an *in vitro* ubiquitylation screen on commercial protein microarrays (ProtoArray®v5.0, ThermoFischer Scientific) containing more than 9000 human proteins expressed and purified as GST-fusion proteins from insect cells. Conditions that we have previously used for *in vitro* ubiquitylation of the SCF^Fbxw5^ substrate Eps8 served as a blueprint for the screen^24^. Two arrays were incubated with recombinant E1, E2s (UbcH5b and Cdc34 together), Ubiquitin, neddylated SCF^Fbxw5^ complex (generated by a split and co-express method^43^) and an ATP regenerating system to ensure reproducibility (Fig 1A). As a control, two arrays were probed with the same mix lacking SCF complexes. To detect ubiquitylated proteins, arrays were stringently washed, incubated with FK2 antibodies specific for conjugated ubiquitin^44^, scanned, quantified and normalised (Fig 1B). As shown in Fig EV1A duplicate reactions were highly reproducible. SCF^Fbxw5^-specific ubiquitylation substrates were then selected based on two criteria: first, signal intensity above an arbitrary threshold (i.e. 500 AU) and second, a more than 5 fold increase in signal intensity over control arrays lacking E3 ligase (Fig 1C and Fig EV1B). We identified a total of 161 candidates (Supplementary Table 1) fulfilling the above-mentioned criteria. Among them were the previously identified substrates Sas6 and Sec23b, demonstrating the capability of the *in vitro* system to identify SCF^Fbxw5^ targets (Fig 1C, note that Ask1 and Eps8 were not present on the array). Gene ontology enrichment analysis of cellular components via the DAVID webtool^45,46^ revealed that a high proportion of these substrates localise to the cytoplasm, which is in line with the described cytoplasmic localisation of Fbxw5^25^. Furthermore, components of the cytoskeleton and vesicle-based transport were enriched among the SCF^Fbxw5^ substrates (Fig 1D). This fits well to the previously identified substrates Eps8, Sas8 and Sec23b and suggests that Fbxw5 could act as a master regulator of cytoskeletal-dependent transport processes. In order to further validate our screen, we picked candidates from these categories and tested their ubiquitylation efficiency by SCF^Fbxw5^ in solution with purified substrates. All selected candidates were efficiently ubiquitylated (Fig 1E) and a high proportion of these substrates interacted with Fbxw5 in co-immunoprecipitation experiments (Fig EV1C and D), demonstrating the high reliability of our screen.

**Figure 1:**
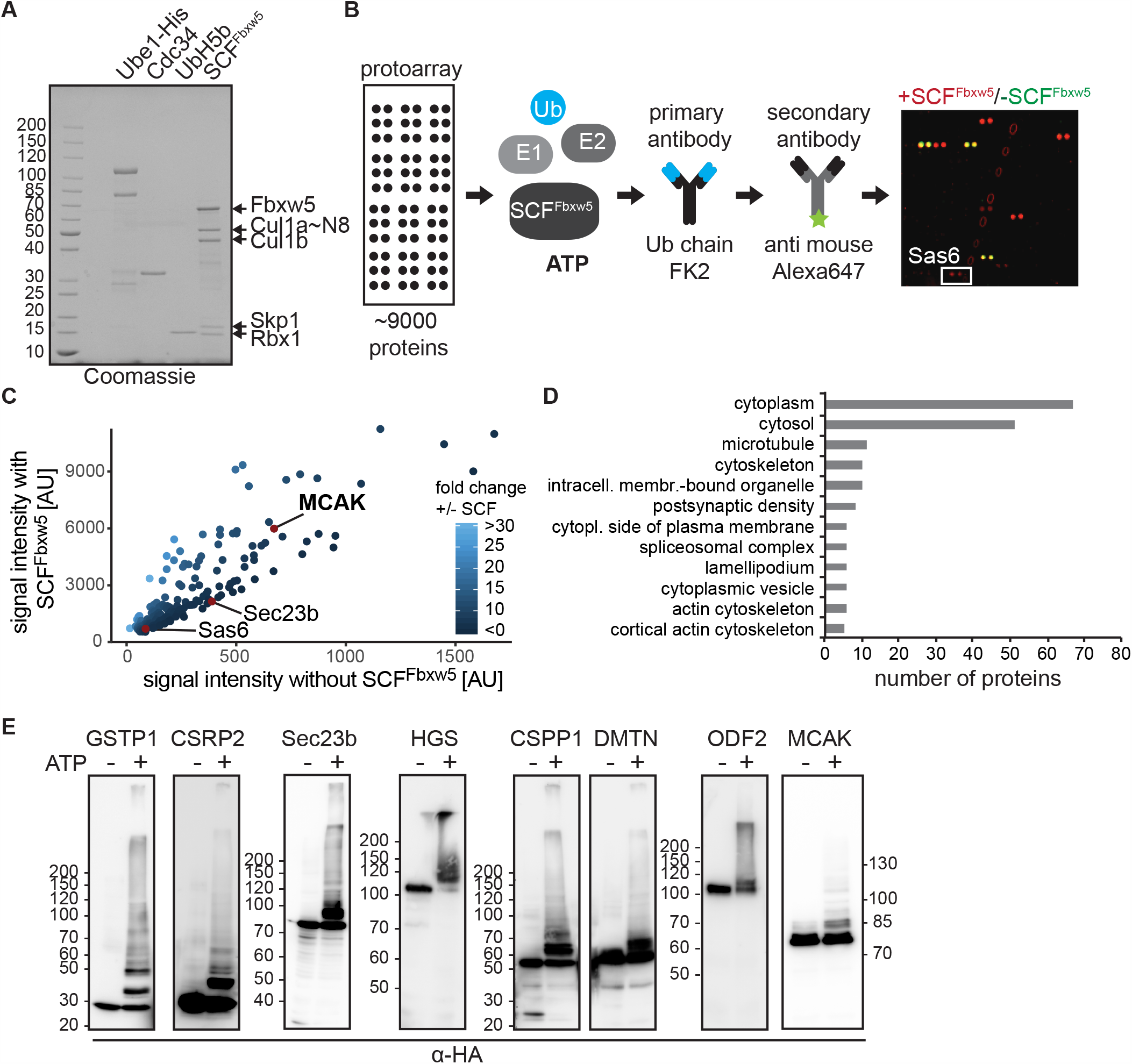
Comprehensive substrate screening on protein microarrays (protoarray®) identifies 161 candidate substrates of SCF^Fbxw5^. **A**. Coomassie-stained SDS-polyacrylamide gel electrophoresis (PAGE) of different proteins used in the assay. Cul1 was obtained using a split and co-express system followed by *in vitro* neddylation. SCF^Fbxw5^ complexes were prepared by mixing equimolar amounts of Fbxw5/Skp1 and Cul1∼Nedd8/Rbx1 sub-complexes. Numbers left of the gel indicate molecular weight marker in kilo-Dalton (kDa, same accounts for all following gels and blots). **B**. Workflow and example of a sub-array of the protoarray screen. Protein microarrays containing more than 9000 human proteins spotted in duplicates were incubated with 15 µM FITC-labelled ubiquitin (Ub), 100 nM E1 (Uba1-His_6_), E2s (0.5 µM each of UbcH5b and Cdc34) and 0.15 µM SCF^Fbxw5^ for 1.5 hours at 37°C. Right panel shows overlay of a selected sub-array probed with (red) or without (green) SCF^Fbxw5^ complexes. White box marks the established substrate Sas6. **C**. Comparison of protoarray signal intensities of candidate substrates probed with or without SCF^Fbxw5^ complexes. Sas6, Sec23b and MCAK are marked as red dots (other published substrates (e.g. Ask1, Eps8) were not among the 9000 proteins spotted on the array). Note that axes have different scaling. **D**. Cellular components GO analysis of identified substrates using DAVID webtool with protoarray proteins as background. **E**. Validation of individual targets by manually curated *in vitro* ubiquitylation experiments. HA-tagged (hemagglutinin) candidate proteins were purified from Hek293T cells via anti-HA immunoprecipitation followed by HA-peptide elution. Candidates were incubated with 20 µM His_6_-Ubiquitin, 170 nM E1, E2s (0.5 µM each of UbcH5b and Cdc34) and 5 mM ATP in the presence or absence of 0.1 µM SCF^Fbxw5^ for 2 hours at 37°C. Substrates were detected via SDS-PAGE followed by western blotting using anti-HA antibodies for detection. Source data are presented in Dataset EV1.

### Fbxw5 interacts with kinesin-13 family members

Due to its pivotal role in different cellular processes and its strong signal intensity in the screen, we focused on the candidate Kif2C/MCAK for further studies. Since ubiquitylation by SCF complexes requires substrate recruitment, we first tested if MCAK interacts with the substrate receptor Fbxw5. As shown by co-immunoprecipitation of tagged proteins, MCAK binds to full length and an F-box lacking mutant of Fbxw5, but not to Fbxw7, indicating an interaction that is independent of other SCF components (Fig 2A). Accordingly, *in vitro* pull-down experiments using purified proteins mixed with competing *E*.*coli* lysates revealed stoichiometric precipitation of MCAK with the Fbxw5/Skp1 sub-complex with no other specific proteins present in the pull-down, confirming that MCAK binds Fbxw5 in a direct and efficient manner (Fig 2B). In order to test if MCAK binds Fbxw5 also in intact cells, we carried out NanoBRET^47^ experiments by overexpressing HaloTag(HT)-tagged Fbxw5 and MCAK fused to NanoLuc luciferase in HeLa cells. Here, Fbxw5 and MCAK generated significantly stronger signals than the negative controls (Fig 2C), confirming that both proteins are also able to interact in living cells. Human cells express two orthologs of MCAK – Kif2A and Kif2B – that have some overlapping and non-overlapping functions in cells^48,49^. Kif2A was present on the protoarray, but was below our threshold and Kif2B was not present on the array. Nevertheless, all three proteins share high sequence similarity and we therefore tested if these orthologs also interact with Fbxw5. Indeed, Kif2B displayed a similar interaction with Fbxw5 in co-IPs and Kif2A precipitated weakly in these experiments (Fig 2D). However it did bind efficiently to Fbxw5 in *in vitro* pull-down assays (Fig 2E). Taken together, our results demonstrate that Fbxw5 can directly recruit all three kinesin-13 proteins.

**Figure 2:**
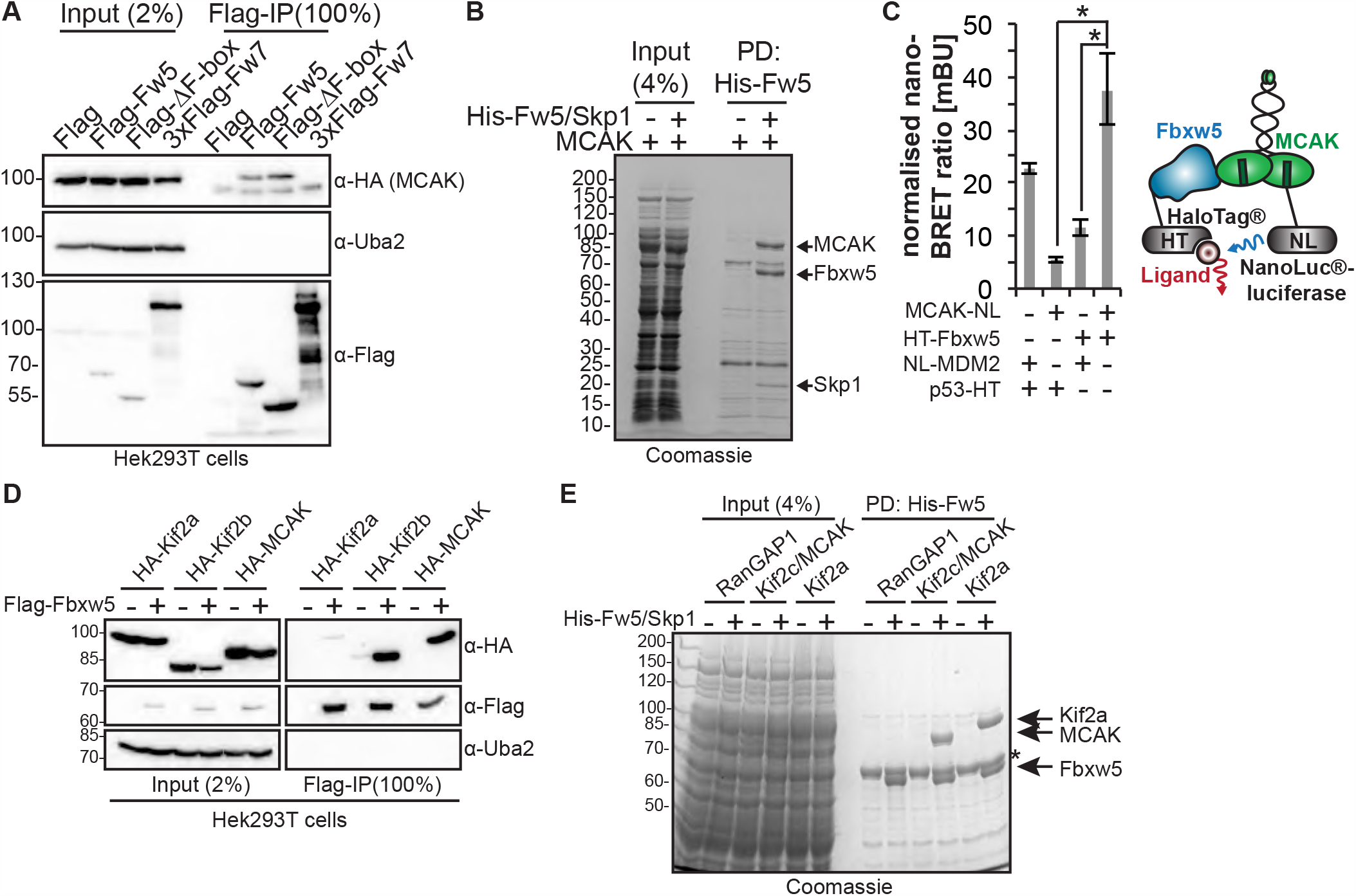
Fbxw5 interacts with kinesin-13 family members. **A**. Co-immunoprecipitation (IP) of HA-MCAK with Flag-Fbxw5. Indicated proteins were expressed in Hek293T cells, extracted and subjected to anti-Flag immunoprecipitation followed by western blot analysis. **B**. *In vitro* pull-down experiment. Indicated proteins (purified from Sf21 cells, 5-fold molar excess of MCAK over Fbxw5/Skp1) were mixed with competing *E*.*coli* lysate and precipitated via Ni-NTA agarose. Proteins were washed, eluted in SDS sample buffer and analysed by SDS-PAGE. **C**. NanoBRET™ (Nano-<u>b</u>ioluminescence <u>r</u>esonance <u>e</u>nergy transfer) assay. Mdm2 and p53 serve as controls. Mdm2 or MCAK were tagged with NanoLuc®-luciferase (Nluc), p53 and Fbxw5 with HaloTag® (HT) and expressed in HeLa cells. After incubation with ligand over night, substrate was added and plates were directly imaged. Left: Quantification of 3 independent experiments. Error bars indicate standard deviation, asterisks the p-value of a two-tailed unpaired Student’s t-test comparing each control with the MCAK/Fbxw5 pair (* < 0.05, ** < 0.01, *** < 0.001, same for all following quantifications). Right: Cartoon showing the NanoBRET™ principal. **D**. Co-immunoprecipitation of HA-Kif2A, HA-Kif2B or HA-MCAK with Flag-Fbxw5 as in A. **E**. *In vitro* pull-down experiment as in B. Asterisk indicate an unspecific protein from *E*.*coli* Note: Kif2B purification from Sf21 cells was much less efficient and it was therefore not included in the experiment. Source data for C are presented in Source Data Table 1.

### SCF^Fbxw5^ ubiquitylates MCAK in a highly efficient and specific manner

Considering that MCAK was a strong hit in our protoarray screen, we were surprised by its rather modest ubiquitylation efficiency within our validation experiments (Fig 1E). In order to investigate the ubiquitylation reaction in more detail, we generated neddylated SCF complexes in high amounts using the baculoviral expression system (Fig 3A and Fig EV2A and B). One difference between the screen and the control experiments was the nature of ubiquitin (FITC-labelled vs His_6_-tagged). We thus wondered if the His_6_-tag on ubiquitin had any impact on the ubiquitylation efficiency of MCAK. Indeed, while MCAK was only weakly ubiquitylated with His_6_-tagged ubiquitin, the reaction became much more accelerated using untagged ubiquitin (Fig EV2C). Of note, ubiquitylation efficiency of Eps8 was not impaired by the His_6_-tag (Fig EV2D). In addition, the effect was specific for the His_6_-tag, as MCAK could be efficiently ubiquitylated with Flag-tagged ubiquitin (Fig EV2E). One explanation for this observation may be a higher positive net charge on the surface of MCAK that repels the His_6_-tag on ubiquitin, but it shows in general that care must be taken when using tags in ubiquitiylation experiments.

**Figure 3:**
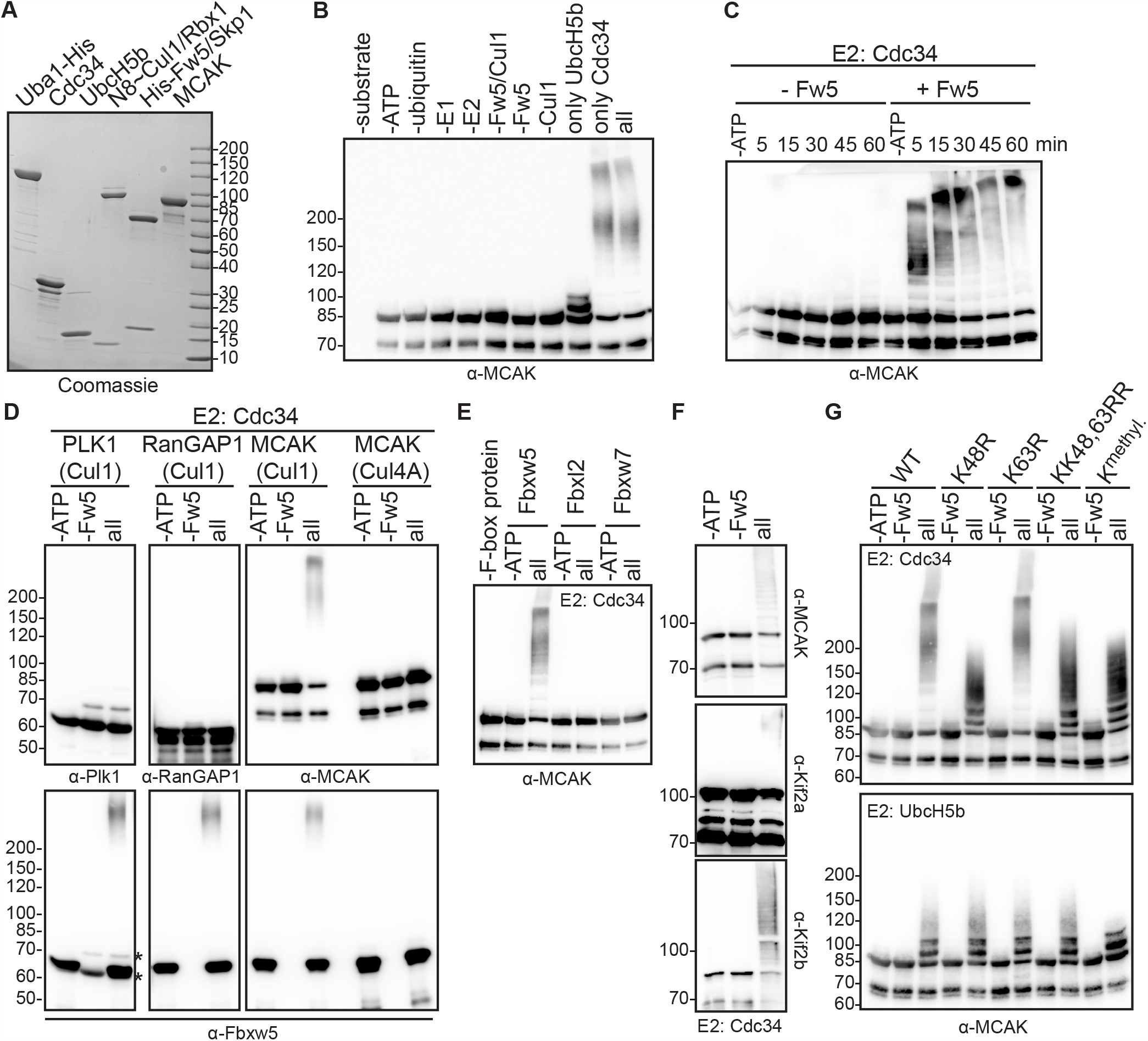
SCF^Fbxw5^ ubiquitylates MCAK in a highly efficient and specific manner. **A**. Coomassie-stained SDS-PAGE of different proteins used in the *in vitro* ubiquitylation assay. Cul1 was obtained as a full-length protein from Sf21 cells and *in vitro* neddylated (see Fig EV2A and B). SCF^Fbxw5^ complexes for all ubiquitylation experiments were prepared by mixing equimolar amounts of Fbxw5/Skp1 and Cul1∼Nedd8/Rbx1 sub-complexes. **B**. Drop-out ubiquitylation experiment. 0.2 µM MCAK was incubated with 75 µM ubiquitin (untagged), 170 nM E1, 0.25 µM of UbcH5b and/or Cdc34 and 0.05 µM SCF^Fbxw5^ for 15 min at 30°C, followed by western blotting using anti-MCAK antibodies for detection. Top labelling indicates component(s) that have been omitted from the reaction. **C**. *in vitro* ubiquitylation experiment as in B using Cdc34 as E2, stopped at the indicated time points by adding SDS sample buffer. **D**. *in vitro* ubiquitylation experiment as in B. Blots 1&2: Plk1 or RanGAP1 used as substrates instead of MCAK. Blot 3: Left: Experiment as in Asterisks indicate signals from Plk1 antibody (same species as anti-Fbxw5) Right: Same reaction except that Cul1∼Nedd8/Rbx1 complexes were replaced by Cul4A∼Nedd8/DDB1/Rbx1 complexes (see Fig EV2A and B). Bottom blots: Same blots as above incubated with anti-Fbxw5 antibodies. Autoubiquitylation of Fbxw5 indicates activity of the E3 ligase complex. **E**. *in vitro* ubiquitylation experiment as in B except that different F-box/Skp1 sub-complexes (Fbxw5/Skp1, Fbxl2/Skp1 or Fbxw7/Skp1) were used (see Fig EV2h). **F**. *in vitro* ubiquitylation experiment as in B using *E*.*coli*-derived MCAK, Kif2A or Kif2B as substrates and Cdc34 as E2. **G**. *in vitro* ubiquitylation experiment as in B using either wild type, K48R, K63R, KK48,63RR or methylated ubiquitin (K^methyl^, see Fig EV2k). Top: Cdc34 used as E2. Bottom: UbcH5b used as E2.

Using untagged ubiquitin for further characterisation, we could confirm that the slower migrating bands of MCAK in the reaction are due to modification by ubiquitin, as they were absent upon drop out of any essential component of the ubiquitin system (Fig 3B). Furthermore, we found that the reaction was much more pronounced using Cdc34 as an E2 compared to UbcH5b and combination of both did not further improve the reaction (Fig 3B and C and Fig EV2F). Unrelated proteins like Plk1 or RanGAP1 were not ubiquitylated (Fig 3D) and the reaction was highly specific to SCF^Fbxw5^ as either replacing Cul1/Rbx1 with Cul4A/DDB1/Rbx1 (Fig 3D and Fig EV2A, B and G) or Fbxw5 with other F-box proteins such as Fbxl2 or Fbxw7 (Fig 3E and Fig EV2H and I) completely abolished MCAK ubiquitylation. Consistent with our previous findings on Eps8, *E*.*coli*-derived MCAK was also ubiquitylated (Fig EV2J), demonstrating that the reaction does not require preceding post-translational modifications on the substrate. In line with the observed binding capability, *E*.*coli*-derived Kif2A and Kif2B were also efficiently ubiquitylated (Fig 3F), showing that SCF^Fbxw5^ is able to target all three orthologs.

Since ubiquitin can form chains of different linkage types with distinct physiological outcomes, we tested whether one of the two most prevalent ones (i.e. K48 and K63) are catalysed by SCF^Fbxw5^ on MCAK. Using UbcH5b as an E2, the ubiquitylation pattern of MCAK was not affected by any of the mutants including methylated ubiquitin that lacks functional acceptor amino-groups (Fig 3G and Fig EV2K). This suggests that UbcH5b mainly catalyses mono-ubiquitylation on MCAK, as it has been shown also for the SCF^βTrCP^ substrate IκBα^50^. Reactions using Cdc34 as E2 on the other hand showed massive loss of high molecular weight species of MCAK and Kif2A using either the K48R single or KK48,63RR double mutant, but not with K63R ubiquitin (Fig 3G and Fig EV2l). The pattern of K48R was the same as for methylated ubiquitin and displayed high similarity with the one obtained with UbcH5b, demonstrating that SCF^Fbxw5^ in concert with Cdc34 catalyses almost exclusively K48 chains on several lysine residues of MCAK and Kif2A.

### SCF^Fbxw5^ affects centrosomal MCAK levels in G_0_

Since K48 chains provide an efficient signal for proteasomal degradation, we examined MCAK levels in RPE-1 cells upon Fbxw5 knockdown. Compared to non-targeting siRNA, MCAK amounts were slightly increased in asynchronously growing cells. Similar effects could be observed after G_2_ arrest using the Cdk1 inhibitor RO-3306^51^ and upon mitotic arrest using nocodazole, in which MCAK levels were in general higher^52^ (Fig 4A and B). However, the most pronounced difference appeared in quiescent cells that had been serum-starved for 24 hours. Here, MCAK levels went almost below detection in control samples, but were 4-fold higher upon Fbxw5 knockdown. Due to this strong effect, we focused on serum-starved cells and used immunofluorescence to test if a specific sub-population of MCAK becomes increased after Fbxw5 knockdown. We detected one prominent signal that co-localised with centrosomes (shown by ODF2 staining) and proved to be specific as it disappeared after MCAK knockdown (Fig 4C). In line with results from western blotting, this signal was on average significantly stronger after Fbxw5 knockdown using three different siRNAs, suggesting that Fbxw5 affects centrosomal MCAK levels upon serum starvation.

**Figure 4:**
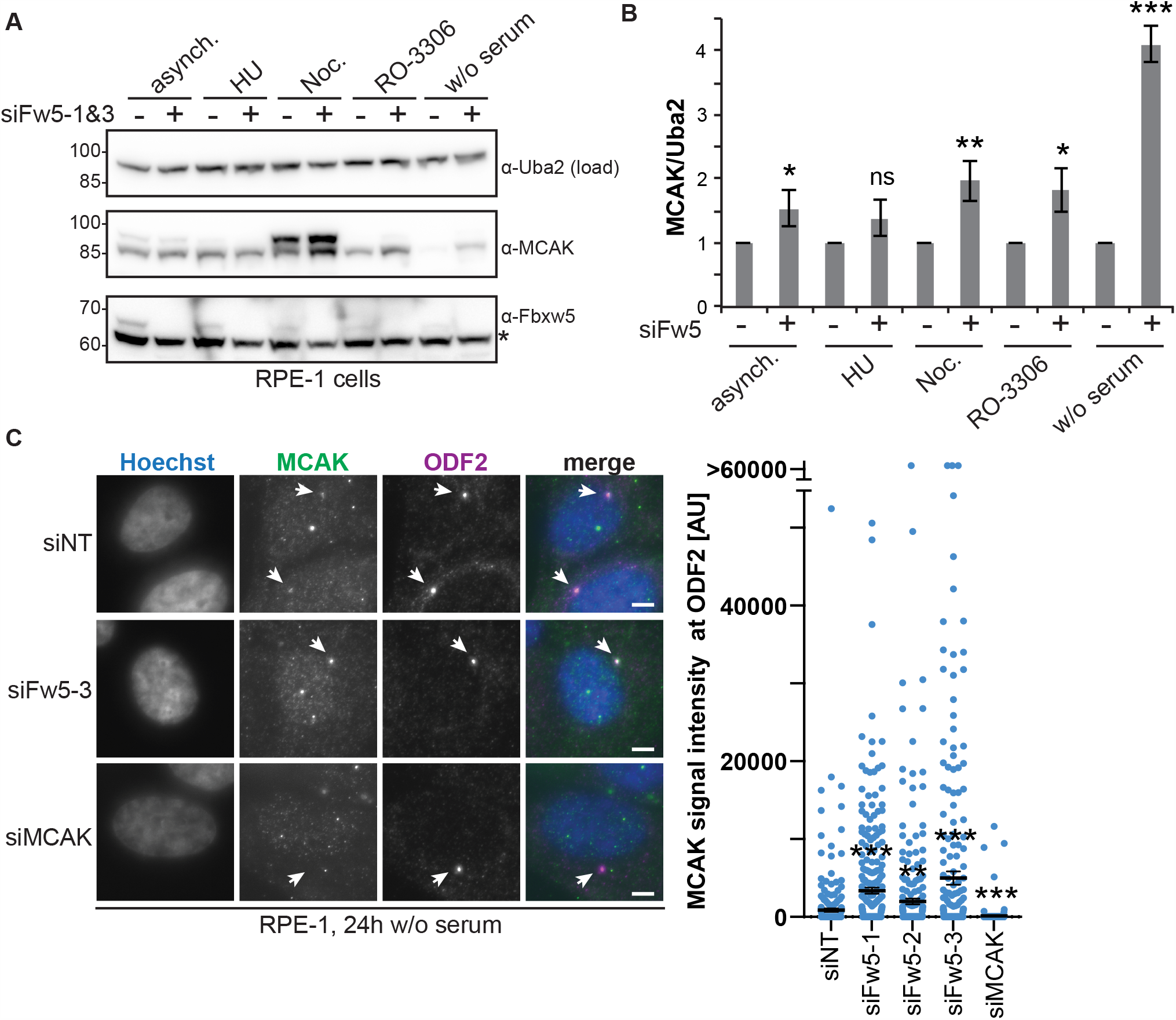
SCF^Fbxw5^ affects centrosomal MCAK levels in G_0_. **A**. Extracts of RPE-1 cells treated for 72 hours with either non-targeting or Fbxw5-directed siRNAs (mix of siFw5-1&3) either grown asynchronously or arrested using 2 mM HU (S-phase), 75 ng/ml nocodazole (M), 10 µM RO-3306 (G_2_) for 18 hours or via serum-starvation for 24 hours (G_0_). Indicated antibodies were used for detection. Asterisk indicates an unspecific band detected by the Fbxw5 antibody. Uba2 served as a loading control (same for all following blots). **B**. Quantification of the MCAK/Uba2 signal ratio normalised to each non-targeting control of 4 independent experiments. Error bars indicate standard deviation, asterisks the p-value of a Student’s t-test comparing each Fbxw5-directed siRNA sample with the non-targeting control. **C**. RPE-1 cells were transfected with the indicated siRNAs (72 hours total time), split on coverslips 24 hours later and serum-starved for the last 24 hours. Cells were fixed in methanol and analysed via immunofluorescence using the indicated antibodies together with Hoechst staining. Left: Maximum intensity projections of representative images. Arrows indicate ODF2 signal. Scale bar = 5 µm. Note: Middle panel shows an example image of siFw5-3 (examples of siFw5-1 and siFw5-2 are not shown). Right: Quantification of MCAK signals co-localising with ODF2. Long black line shows mean intensity and error bars indicate standard error of the mean of 3 independent experiments covering in total more than 280 cells. Asterisks indicate p-value of a Mann Whitney test comparing each sample set with the non-targeting control. Source data for B and C are presented in Source Data Table 1.

### SCF^Fbxw5^ and the APC/C regulate MCAK at distinct cell cycle stages

Considering the strong difference in MCAK levels upon serum starvation, we initially hypothesised that the Fbxw5-dependent regulation occurs upon mitotic exit when cells enter into a quiescent state. To test this, we generated a stable cell line expressing mNeonGreen-(mNG-)tagged MCAK under a doxycyclin-inducible promoter. As shown in Fig EV3A mNG-MCAK localisation was very similar to endogenous MCAK observed by IF. We then used these cells to perform fluorescence-based pulse-chase experiments. For this, we transfected cells with the according siRNAs, and induced mNG-MCAK expression 24 hours later. After another 24 hours we removed doxycyclin and released cells into serum-free medium. Using spinning disk microscopy, we then selected cells in telophase and imaged them for 20 hours with 20 minutes time intervals (Fig 5A and Movie EV1). To our surprise, mNG-MCAK signal gradually disappeared both in control and Fbxw5 knockdown cells with almost equal half-life times for the centrosomal population. Similar results were obtained for endogenous MCAK in an IF-based mitotic shake off experiment, in which MCAK levels became gradually fainter at centrosomes in both control and knockdown cells (Fig EV3B).

**Figure 5:**
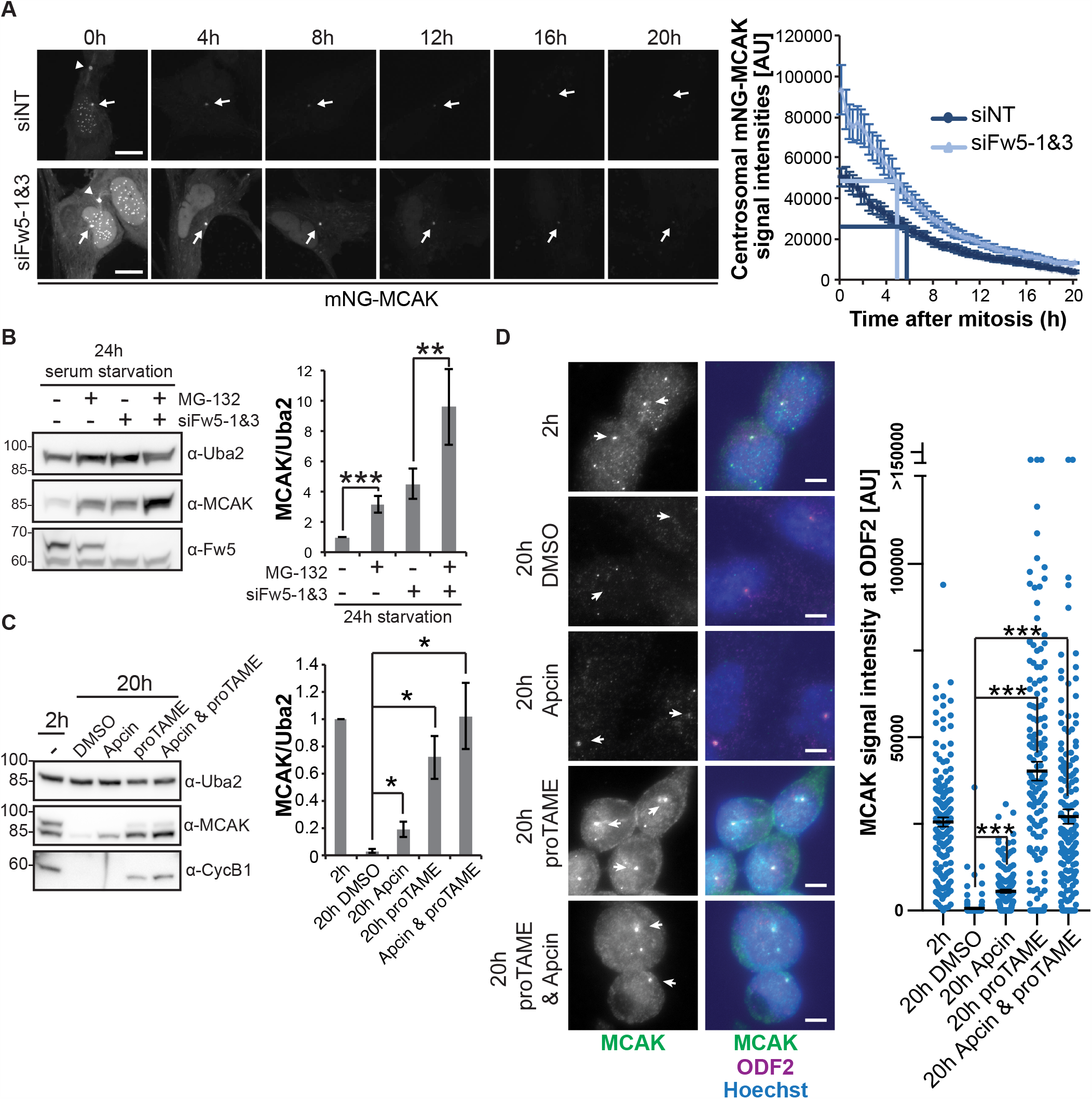
The APC/C regulates MCAK after mitosis. **A**. Fluorescence-based pulse-chase experiment using a monoclonal RPE-1 cell line expressing N-terminally mNeonGreen-(mNG)-tagged MCAK under a doxycyclin-inducible promoter. Cells were transfected with the indicated siRNAs (siFw5-1&3 together) and split 24 hours later on Ibidi 8 Well Glass Bottom µ-slides while simultaneously inducing mNG-MCAK expression with 6 ng/ml doxycyclin (pulse). 24 hours later, doxycyclin was washed out and cells were released into serum-free medium. Cells that have just finished or were about to finish mitosis (distinguishible for example by the midbody signal of mNG-MCAK (arrow heads)) were selected and imaged over 24 hours, taking an image every 20 min (chase). Left: representative images (maximum intensity projections) of selected time points. Scale bar = 10 µm. Arrows indicate centrosomal mNG-MCAK signals. See also Movie EV1. Right: Quantification of centrosomal mNG-MCAK signals. Error bars show standard error of the mean of 3 independent experiments with 25 cells in total. P-value of a Students t-test comparing signal intensity of Fbxw5 knockdown over non-targeting control was mostly below 0.01 for time points 0 to 8 hours and below 0.05 for time points 9h to 20h. **B**. Extracts of RPE-1 cells treated with the indicated siRNA for 72 hours, serum starved for the last 24 hours and treated either with DMSO (-) or 10 µM MG-132 (+) for the very last 6 hours. Left: Representative image. Right: Quantification of MCAK/Uba2 signal ratio normalised to non-targeting control with DMSO of 5 independent experiments. Error bars indicate standard deviation, asterisks the p-value of a Student’s t-test as above. **C**. RPE-1 cells were treated with 75 ng/ml nocodazole for 4 hours. Mitotic cells were shaken off, washed 2 times with phosphate-buffered saline (PBS) and released in serum free medium. After 2 hours, cells were either directly harvested (2 hour sample) or treated with the indicated APC/C inhibitor for another 18 hours before harvesting. Left: Representative blot. Right: Quantification. Error bars show standard deviation of 3 independent experiments, asterisks the p-value of a Student’s t-test as above. **D**. Same samples as in C, except that cells grown on coverslips were fixed in methanol and analysed by immunofluorescence. Left: Maximum intensity projections of representative images. Arrows indicate ODF2 signal. Scale bar = 5 µm. Right: Quantification. Error bars show standard error of the mean of 3 independent experiments, covering in total more than 150 cells. Asterisks indicate the p-value of a Mann Whitney test. Source data are presented in Source Data Table 1.

The observation that endogenous and mNG-tagged MCAK slowly disappear upon entry into a quiescent G_0_ state in an Fbxw5-independent manner indicates the existence of another regulatory pathway. In line with this, proteasomal inhibition via MG-132 during the last 6 hours of serum starvation significantly increased total MCAK levels in control but also in Fbxw5 knockdown cells (Fig 5B). One likely candidate for an alternative regulatory pathway is the Anaphase-Promoting Complex/Cyclosome (APC/C) – another multi-subunit ubiquitin E3 ligase that has been shown to target MCAK via Cdc20 in HeLa cells^53^ and via Cdh1 *in vitro*^54,55^. In order to test if the APC/C targets MCAK after mitotic exit, we again performed mitotic shake-off experiments with release into serum-free medium. This time, we added the APC/C inhibitors Apcin and proTAME after mitotic completion (∼2 hours after shake-off and nocodazole washout). While Apcin prevents substrate recognition by masking the D-Box binding site of Cdc20, proTAME inhibits incorporation of both substrate receptors (Cdc20 and Cdh1) into the APC/C complex. Combination of both compounds has been shown to completely block APC/C activity^56,57^. As expected, total and centrosomal MCAK signals disappeared almost completely 20 hours after mitosis in DMSO-treated samples (Fig 5C and D). However, this effect was partially reverted in Apcin-treated cells and almost completely reverted upon proTAME addition, suggesting that the APC/C is indeed responsible for MCAK removal upon entry into quiescence. Taken together, our findings show that after mitotic exit MCAK is predominantly regulated by the APC/C.

### SCF^Fbxw5^ regulates MCAK in G_2_/M

The experiments described above did not reveal any stabilisatory effect on MCAK upon Fbxw5 depletion after mitotic exit. However, one striking difference that was revealed in the time course experiments (Fig 5A and Fig EV3B) were the elevated levels of MCAK in Fbxw5 knockdown cells at time point zero. Accordingly, depletion of Fbxw5 led to elevated mNG-MCAK levels already in prometaphase (Fig EV3C). As mNG-MCAK expression was induced only for 24 hours in this experiment, these results imply that Fbxw5 targets MCAK before mitotic exit. Since we were not able to detect ubiquitylated species of MCAK using different enrichment procedures such as TUBES^58^ or FK2 antibodies (data not shown), we employed cycloheximide (CHX) chase experiments in different pre-mitotic cell cycle arrests to further narrow down the exact timing of the Fbxw5-dependent regulation. Whereas MCAK remained relatively stable for 6 hours in asynchronously growing and in S phase arrested cells, it became unstable in cells arrested in G_2_ with the Cdk1 inhibitor RO-3306 in an Fbxw5-dependent manner (Fig 6A). This is in line with our previous work on Eps8, which was also targeted during G_2_^24^, indicating that SCF^Fbxw5^ is particularly active during this cell cycle stage.

**Figure 6:**
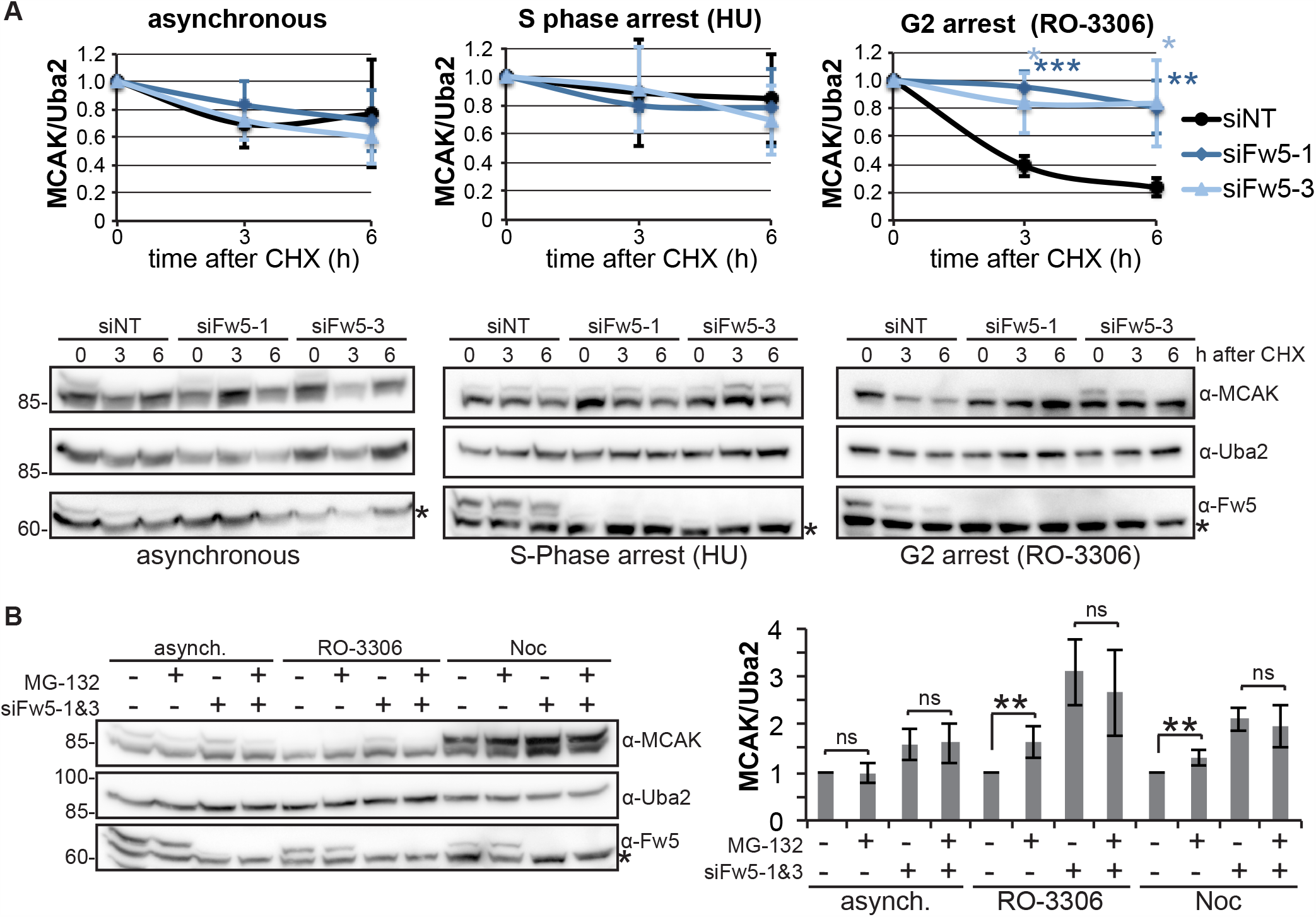
SCF^Fbxw5^ regulates MCAK in G_2_/M. **A**. Cycloheximide (CHX) chase experiments of RPE-1 cells treated with the indicated siRNA for 48 hours and then either grown further asynchronously or arrested at S phase with 2 mM hydroxyurea (HU) or at G_2_ with 10 µM RO-3306 for 18 hours. 50 µg/ml CHX was added to the cells, samples were harvested at the indicated time points and analysed via SDS-PAGE and western blotting using the indicated antibodies for detection. Bottom: Representative images. Asterisks indicate an unspecific band detected by the Fbxw5 antibody. Top: Quantification of MCAK/Uba2 signal intensity ratios normalised to time point zero. Error bars show standard deviation of 4 independent experiments and asterisks indicate p-value of a Student’s t-test. **B**. Extracts of RPE-1 cells treated with the indicated siRNAs for 72 hours, grown asynchronously or arrested with the indicated compounds for the last 24 hours and treated either with DMSO (-) or 10 µM MG-132 (+) for the very last 6 hours. Left: Representative image. Asterisk indicates an unspecific band detected by the Fbxw5 antibody. Right: Quantification of MCAK/Uba2 signal ratio normalised to non-targeting control with DMSO of 6 independent experiments. Error bars indicate standard deviation, asterisks the p-value of a Student’s t-test as described above. Source data for A and B are presented in Source Data Table 1.

To gain further evidence that Fbxw5 targets MCAK specifically during G_2_, we investigated MCAK amounts upon proteasomal inhibition. In contrast to asynchronously growing cells, MG-132 treatment under G_2_ arrest led to a moderate but reproducible increase in MCAK amounts (Fig 6B). Since MG-132 treatment did not further increase MCAK levels upon Fbxw5 knockdown, these results confirm that MCAK becomes proteasomally degraded in an Fbxw5-dependent manner during G_2_. Similar results were obtained for nocodazole-arrested cells. However, due to the incompatibility of CHX with mitotic arrested cells^59^, we were not able to further confirm this effect. Nevertheless, our results together demonstrate that the activity of SCF^Fbxw5^ towards MCAK starts during G_2_, probably persist during mitosis and becomes undetectable after mitotic exit.

### Fbxw5-dependent regulation of MCAK in G_2_/M is required for ciliogenesis

Since excess activity of MCAK has been shown to increase mitotic duration^35,60^, we speculated that its regulation by Fbxw5 in G_2_/M may be an essential process for mitosis. However, we did not observe a striking increase in the mitotic index of RPE-1 cells (Fig EV4A) and mild effects such as a delay in prometaphase may well be due to other Fbxw5 substrates like Sas6 or Eps8, which have been shown to impact on mitotic progression, too^24,25^.

A second period during which elevated MCAK levels could be detrimental is the G_0_ phase. In fact, overexpression of kinesin-13 proteins (including MCAK) has recently been demonstrated to impair ciliogenesis in RPE-1 cells upon serum starvation^42^. Consistent with this, we observed a dramatic decrease in ciliated cells upon MCAK overexpression in serum-starved RPE-1 cells using our doxycyclin-inducible system. Interestingly, this defect was dependent on the timing of MCAK overexpression relative to serum withdrawal. Whereas inducing MCAK overexpression before serum starvation provoked a massive loss of ciliated cells, it had only little impact when initiated in cells that had been already serum starved for 24 hours (Fig 7A). Regardless of the timing, cilia that were still present upon MCAK overexpression showed a significant reduction in their length that correlated with MCAK intensities at basal bodies (Fig EV4B and C). Taken together, these results suggest that excess centrosomal MCAK strongly inhibits the de novo generation of cilia, but only decreases the lengths of axonemes on pre-existing ones without leading to a complete resorption in most cases.

**Figure 7:**
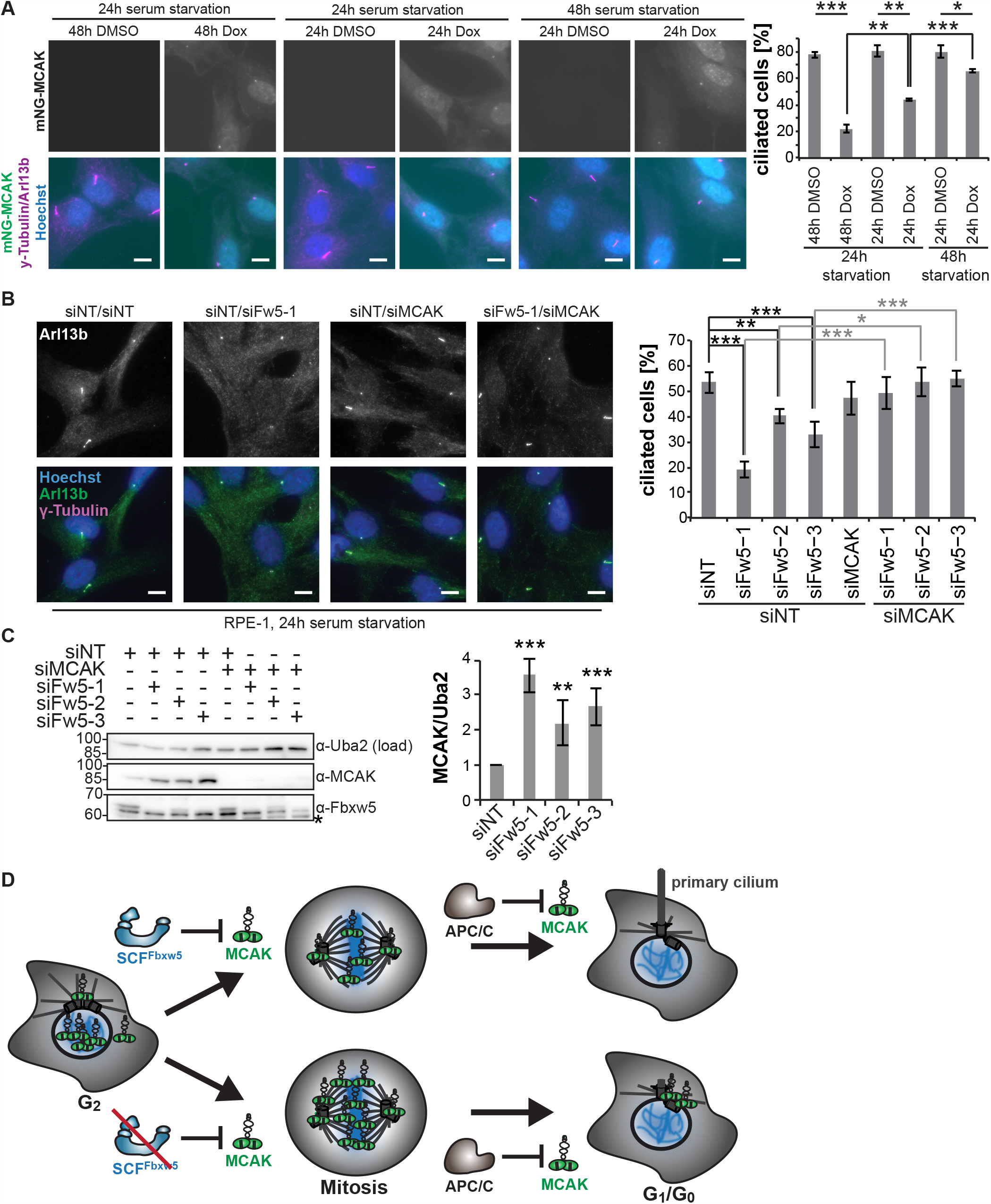
Fbxw5 is required for ciliogenesis in an MCAK-dependent manner. **A**. Polyclonal RPE-1 cells expressing mNG-MCAK under a doxycyclin-inducible promoter were seeded on coverslips and grown for 24 hours. Subsequently, cells were washed and incubated for another 24 or 48 hours in serum-free medium. During this procedure, mNG-MCAK expression was induced with 20 ng/ml doxycycline either 24 hours before, concomitantly with, or 24 hours after serum withdrawal. Cells were then fixed in methanol and subjected to immunofluorescence using the indicated antibodies. Left: Maximum intensity projections of representative images. Scale bar = 10 µm. Right: Quantification of ciliated cells. Error bars show standard deviation of 3 independent experiments covering more than 200 cells and asterisks indicate p-value of a Student’s t-test. **B**. RPE-1 cells were treated with the indicated siRNA for 48 hours followed by serum-starvation for another 24 hours. Cells were then fixed in methanol and subjected to immunofluorescence using the indicated antibodies. Left: Maximum intensity projections of representative images. Note: Left panel shows an example image of siFw5-1 (examples of siFw5-2 and siFw5-3 are not shown). Scale bar = 10 µm. Right: Quantification of ciliated cells. Error bars show standard deviation of 4 independent experiments covering more than 200 cells and asterisks indicate p-value of a Student’s t-test. **C**. Western blot of RPE-1 cell extracts. Left: Representative image. Asterisk indicates an unspecific band detected by the Fbxw5 antibody. Right: Quantification of MCAK/Uba2 signal ratio. Error bars show standard deviation of 6 independent experiments and asterisks the p-value of a Student’s t-test comparing each knockdown with the non-targeting control. **D**. Model: MCAK is regulated by two distinct E3 ligases at different time points. While SCF^Fbxw5^ targets MCAK for degradation during G_2_/M, the APC/C takes over after mitotic exit. Via the SCF^Fbxw5^ pathway, cells ensure that MCAK levels are kept low upon entry into G_0_ and thus permit ciliogenesis in the following cell cycle. Source data for A, B and C are presented in Source Data Table 1.

Since loss of Fbxw5 increased MCAK levels at centrosomes during serum starvation (Fig 4C), we wondered if Fbxw5 knockdown induces similar defects in ciliogenesis of wild type RPE-1 cells. Indeed, knockdown of Fbxw5 with three different siRNAs led to a significant reduction in ciliated cells (Fig 7B). Similarly to MCAK overexpression, remaining cilia under these conditions displayed on average much shorter axonemes (Fig EV5D). Strikingly, simultaneous knockdown of MCAK almost completely rescued both Fbxw5-dependent ciliogenesis phenotypes, indicating that these effects are due to elevated levels of MCAK and not of any other substrate. In conclusion, our data demonstrate that loss of Fbxw5 impairs ciliogenesis via elevated MCAK levels at the beginning of G_0_.

## Discussion

Using comprehensive substrate screening on protein microarrays, we identified MCAK as a bona fide substrate of SCF^Fbxw5^, assigned its regulation pathway to the G_2_/M phase of the cell cycle and demonstrated that this process is required for ciliogenesis upon entry into a quiescent state. In addition, our *in vitro* ubiquitylation screening approach provides a useful and reliable source for potential Fbxw5 substrates. The number of 161 candidate substrates may seem high at first glance, but one has to keep in mind that proteins are probed here in a cell-free system. Fbxw5 may regulate some of its substrates only in certain cell types or during specific developmental stages. In contrast to cell-based screens, the protein microarray method is unique in yielding a comprehensive list of potential substrates without being limited to a particular cellular context.

The identification of MCAK and its orthologs as substrates of SCF^Fbxw5^ is an important step towards a better understanding of these important microtubule depolymerases. Besides their well-explored roles in mitosis, MCAK and its orthologs have been identified recently as negative regulators of ciliogenesis^42^. We could show that Kif2A and Kif2B are also ubiquitylated *in vitro* by Fbxw5 and it is therefore possible that stabilisation of these proteins (or other substrates) contributes to the ciliogenesis defect observed upon Fbxw5 loss. However, concomitant knockdown of MCAK almost completely restored cilia formation, suggesting that either MCAK alone is responsible or that lowering the activity of microtubule depolymerases in general is sufficient to regain ciliogenesis.

How MCAK regulates ciliogenesis is still elusive. Besides a reduction of ciliated cells, we also observed a shortening of remaining cilia in Fbxw5 knockdown or MCAK overexpressing cells. Taking into account MCAK’s ability to depolymerise microtubules at both ends^32^, this implies that MCAK could indeed be able to exert its microtubule activity at basal bodies, similarly to what has been proposed for Kif2A^42^. Interestingly, overexpression of MCAK impaired cilia formation to different extends depending on the time point of its induction relative to serum withdrawal. Whereas MCAK overexpression 24 hours after serum starvation only provoked little reduction in ciliated cells, it massively impaired ciliogenesis when induced 24 hours prior to serum withdrawal. Although we observed an increase in mitotic cells by 5% upon MCAK overexpression in full serum (Fig EV4E), this cannot fully explain the massive loss of ciliated cells from 80 to 20%. Instead, we propose that high levels of MCAK become particularly deleterious at initial stages of ciliogenesis. Here, newly synthesised axonemal microtubules may be better accessible for MCAK-dependent depolymerisation due to their shorter length, a potentially incomplete microtubule lattice or a lack of posttranslational modifications that may impair MCAK activity such as detyrosination^61^.

Whereas further work is certainly necessary to determine the exact mode of action, this theory could also explain why Fbxw5 targets MCAK in G_2_/M. Such a preceding regulatory event may be required to guarantee the reduction of MCAK levels at the beginning of the quiescent state in order to allow timely formation of primary cilia. Despite an on-going removal via the APC/C after mitosis, cells lacking Fbxw5 enter G_0_ with much higher MCAK levels. The concomitant delay in ciliogenesis could be detrimental in settings that require fast cilia formation, for example during tissue and organ development. Hence, placing the regulation process towards the end of the preceding cell cycle may ensure timely removal of negative factors enabling quick cilia-dependent sensing of extracellular signals later on.

Could this be a general concept? To the best of our knowledge, such a preceding regulatory process for ciliogenesis has not been described so far. However, degradation of the centriolar protein CP110 by SCF^CyclinF^ has been shown to occur during G_2_/M, where it ensures centrosome homeostasis during mitosis^62,63^. Other studies have established that CP110 also counteracts ciliogenesis by recruiting kinesin-13 Kif24^64,65^. As far as we know, it has not been demonstrated yet whether the SCF^CyclinF^-dependent regulation event is indeed required for ciliogenesis. But if this turns out to be the case, degradation of negative factors by SCF E3 ligases during G_2_/M may represent a common mechanism that allows timely formation of primary cilia in the following G_1_/G_0_ phase. It will be interesting to see, how such preceding regulatory events impact on cilia-dependent developmental programs within multicellular organisms and whether they play a critical role in ciliopathies.

## Author contributions

J.S. designed and performed most experiments and wrote the manuscript. G.H. designed and performed the protoarray screen and wrote this part of the manuscript, S. Hess designed and performed the BRET assays, F. Mikus designed and performed the postmitotic degradation assays (Fig 5A, Fig EV3B), S. Hata generated the RPE-1 TET3G cell line, K.M. purified various proteins and carried out some ubiquitylation assays, K-P.K. designed the BRET assays and F. Melchior guided the project, designed experiments and wrote the manuscript.

## Acknowledgements

This project received funding from the Deutsche Forschungsgemeinschaft (SPP1365, ME 2279/3 to F.M. and DFG KN590/7-1 to K-P.K), the European Union’s Horizon 2020 research and innovation programme under the Marie Skłodowska-Curie grant agreement N° 748315 (UBIMAPS) and the Heidelberg CellNetworks Cluster of Excellence postdoctoral program (to J.S.). We would like to thank Vladimir Benes for providing access to a microarray reader and Brenda Schulman for the generous gift of recombinant neddylated Cul1/Rbx1 and plasmids of neddylation enzymes. We gratefully acknowledge Giorgio Scita, Eric Fischer, Ludger Hengst, Ingrid Hoffmann, Achim Dickmanns, Sima Lev, Shahri Raasi, Elmar Schiebel, Gislene Pereira and Bahtiyar Kurtulmus for plasmids and advise, and thank Holger Lorenz, Christian Hörth and Monika Langlotz for excellent microscopy and FACS support. Finally, we would like to thank Anthony Razov, Hannah Lee, Lisa Mutz, Moritz Dodenhöft for technical support, Annette Flotho and Roman Beloshistov for critical reading of the manuscript and the whole Melchior lab for helpful discussions, reagents and advise.

## Material and Methods

### Cell culture

RPE-1 hTERT cells (ATCC: CRL-4000) were cultured in Dulbecco’s modified Eagle Medium/Nutrient mixture F12 (DMEM/F12) supplemented with 10% filtrated fetal bovine serum (FBS). Hek293T cells (DSMZ: ACC 635 Lot 4) were cultured in DMEM supplemented with 2 mM glutamine and 10% FBS. Hela cells were cultured in DMEM Glutamax™ supplemented with 10% FBS. All cell lines were authenticated at the DKFZ Genomics and Proteomics Core Facility (Heidelberg), regularly tested for mycoplasm contamination, kept constantly below confluency and were cultivated at 37°C with 5% CO_2_ for no longer than 8 weeks (∼20 passages). In order to induce ciliogenesis, cells were washed 2x with phosphate-buffered saline (PBS) and then incubated with DMEM/F12 medium lacking FBS.

### Plasmid transfection

Hek293T cells were transiently transfected 1 day after seeding in 15-cm dishes at a confluency of 50% with 30 µg total plasmid DNA using polyethyleneimine (PEI, 1 mg/ml pH 7.0) at a DNA:PEI ratio of 1:2.5 in serum-free DMEM. Serum supplemented DMEM was added 6 hours after transfection. Cells were harvested 24-48 hours after transfection.

### siRNA transfection

siRNA transfection was performed with Lipofectamine RNAi MAX (Invitrogen) according to the manufacturer’s recommendation for reverse transfection. For a 6-well format, trypsinised cells were mixed with 500 µl OptiMEM (Gibco) containing 4 µl RNAi MAX and 50 pmol total siRNA and medium was added to reach a total volume of 2.5 ml. After ∼6 hours, cells were washed 1x with PBS, replenished with fresh medium and split on the next day. If not othewise mentioned, cells were harvested or analyzed 48h or 72h after siRNA transfection. siRNAs and all other consumables are listed in Appendix Table 2.

### Generation of stable cell lines

For the generation of stable cell lines expressing mNG-MCAK under a doxycycline-inducible promoter, the retroviral transduction system of Clonetech® “Retro-X™ Tet-On® 3G Inducible Expression System” was used according to the manufacturer’s protocol (retroviral gene transfer and expression user manual). Stable cells expressing low levels of the according protein were sorted using a BD FACSMelody™ or BD FACSAria III (Becton Dickinson) after induction with doxycycline for 24 h.

### Insect cell Culture

Sf21 cells were cultured in Sf-900™ II SFM or ExCell® 420 Serum-Free Medium at 130 rpm 27°C and generally kept between 1×10^6^ and 8×10^6^ cells/ml. Baculoviruses for protein expression were generated using the Bac- to-Bac Baculovirus Expression System (Invitrogen) according to the manufacturer’s instructions. For bacmid generation, the according plasmid was transformed into DH10MultiBac chemically competent cells. Bacmids were isolated, tested for integrity by PCR and transfected into Sf21 cells using Cellfectin-II reagent in a 6-well format. 72 hours or 96 hours after transfection, medium of transfected cells was harvested and filtered to generate P1 viral stocks. High-titer P2 stocks were produced by infecting 50 ml of Sf21 cells with the according P1 stock for 72 hours and used for final protein expression by infecting 100-200 ml of Sf21 cells at 2×10^6^ cells/ml for 48 h. All F-box proteins were purified together with Skp1 by co-infecting Sf21 cells with both P2 viral stocks.

### Protoarray *in vitro* ubiquitylation screen

Human ProtoArray® Microarray (v5.0) containing 9483 unique human proteins expressed as GST-fusion in insect cells were purchased from Invitrogen. Arrays were blocked in 4°C cold blocking buffer (50 mM HEPES pH 7.4, 200 mM NaCl, 0.05% Tween-20, 25% glycerol, 10 mg/ml ovalbumin, 5 mM reduced glutathione, 1 mM dithiothreitol (DTT), 1 µg/ml aprotinin/pepstatin/leupeptin) for 1.5 hours. Arrays were then washed with assay buffer (50 mM Tris pH 7.5, 100 mM NaCl, 5 mM MgCl_2_, 0.05% Tween-20, 2 mg/ml ovalbumin, 1 mM DTT, 1 µg/ml aprotinin/pepstatin/leupeptin) for 5 min. For the reaction, 120 µl ubiquitylation reaction mixture (100 nM Uba1, 0.5 µM UbcH5b, 0.5 µM Cdc34, 15 µM FITC-Ubiquitin, 150 nM SCF^Fbxw5^ (containing split-Cul1 from *E*.*coli*^43^), energy regeneration solution (B-10, Boston Biochem) in assay buffer were carefully pipetted directly on the arrays. The slides were covered with cover slips and incubated in a humid chamber for 1.5 hours at 37°C. Reactions were stopped by submerging slides in 5 ml assay buffer and removing the cover slip. Assay buffer was aspirated immediately and slides were washed twice with BSA buffer (PBS (140 mM NaCl, 2.7 mM KCl, 10 mM Na_2_HPO_4_, 1.5 mM KH_2_PO_4_, pH 7.5), 0.1% Tween-20, 1% BSA). Slides were then washed 3 times with 0.5% SDS followed by 3 more washes with BSA buffer. For detection of ubiquitylated proteins, slides were first blocked with antibody blocking buffer (PBS, 0.1% Synthetic Block (Invitrogen)) for 10 min followed by 1.5 hours incubation at 4°C with anti-ubiquitin chain antibody FK2 (1 µg/ml FK2 in antibody blocking buffer). Modified proteins were finally labelled with Alexa-647 labelled anti-mouse antibodies. Slides were washed, spin-dried (200 g for 1 min) and scanned with a Genpix 4000B microarray scanner (Molecular Devices, made available by EMBL Genomics Core Facility). GenePix® Pro v.6 software (Molecular Devices) was used to align protein spots and quantify fluorescence intensity of the spots. Background correction was performed with Protein Prospector analyzer (Invitrogen) followed by normalisation using protein array analyzer package (v1.3.3)^66^ for R (Version 3.2.1.). Proteins were considered SCF^Fbxw5^ dependent ubiquitylation targets if signal intensity was >500 and more than 5-fold increased over arrays incubated without E3 ligases. All candidate spots were manually controlled on the scanned images.

### GO analysis and statistical analysis

GO analysis of cellular components (CC_Direct) was performed using the DAVID Bioinformatics Resources 6.8. SwissProt IDs of candidate proteins were converted to EntrezGene ID before analysis via http://www.uniprot.org. SwissProt IDs of all proteins in the Human ProtoArray® Microarray (v5.0) were converted to EntrezGene ID accordingly and used as background. All statistical analysis was performed using either Microsoft Excel® (for student’s t-test) for Mac (version 14.6.8) or GraphPad Prism v9 for Mac (for Mann Whitney test and Pearson correlation). For western blot quantification, ratio of MCAK signal over Uba2 signal was calculated, normalised to the non-targeting control and statistical significance was assessed using a two-tailed student’s t-test. Due to thresholding, MCAK signal analysis using immunofluorescence yielded many values of zero intensity and thus generated a non-Gaussian distribution. Hence, a Mann-Whitney test was used to estimate statistical significance.

### Recombinant protein purification

All bacteria-derived proteins were expressed in Rosetta (DE3) cells, all insect cell-derived proteins in Sf21 cells. After purification, final protein samples were aliquoted, snap-frozen in liquid N_2_ and stored at −80°C until further use. Expression and purification of RanGAP1, Eps8, Ubiquitin, Uba1, UbcH5b, UbcH5c, Cdc34, APPBP1-Uba3, Nedd8, Ubc12 and Nedd8*Cul1A-Cul1B/Rbx1 was described before^24,67–71^. For Ube2G1 expression, the same procedure as for UbcH5b and Cdc34 was followed. Briefly, proteins were expressed over night in Rosetta (DE3) cells at 16°C with 200 µM IPTG, lysed in 50 mM Tris pH 8, 300 mM NaCl, 1 mM EDTA, 1 mM β-mercaptoethanol, 1 µg/ml aprotinin/pepstatin/leupeptin by passing two times through an Emulsion Flex-C5 microfluidizer. Afterwards, glutathione-sepharose beads were added, extensively washed and proteins were eluted in 50 mM Tris pH 8.8, 150 mM NaCl, 1 mM β-mercaptoethanol and 25 mM glutathione. Finally, GST-tag was cleaved with Prescission protease overnight and removed by molecular sieving using an analytical Superdex 75 10/300 chromatography column (GE Healthcare).

For His-tagged MCAK and Kif2A purified from *E*.*coli*, protein expression was induced overnight at 16°C with 200 µM IPTG and cells were lysed by passing two times through an Emulsion Flex-C5 microfluidizer in buffer A (25 mM Hepes pH 7.4, 300 mM KCl, 2 mM MgCl_2_, 5 mM β-mercaptoethanol, 30 mM imidazole, 50 µM ATP, 1 µg/ml aprotinin/pepstatin/leupeptin and 1 mM PefaBloc (Roche)). Proteins were purified over Ni-NTA, eluted in buffer A (devoid of ATP and PefaBloc but containing 200 mM instead of 30 mM imidazole) and further purified over a Superdex 200 10/300 GL column (GE Healthcare) in buffer B (25 mM Hepes, pH 7.4, 300 mM KCl, 2 mM MgCl2, 1 mM DTT, 1 µg/ml aprotinin/pepstatin/leupeptin). His-tagged Kif2B was generated accordingly, except that lysis was performed in buffer C (50 mM sodium phosphate buffer pH 6.0, 300 mM KCl, 2 mM MgCl_2_, 5% glycerol, 5 mM β-mercaptoethanol, 30 mM imidazole, 0.06% NP-40, 50 µM ATP, 1 µg/ml aprotinin/pepstatin/leupeptin and 1 mM PefaBloc). Proteins were eluted in buffer C devoid of ATP, NP-40 and PefaBloc but containing 200 mM imidazole and further purified over a Superdex 200 10/300 GL column in buffer B.

For MCAK and Kif2A purified from insect cells, 200 ml of Sf21 cells at 2×10^6^ cells/ml were infected with 1% (v/v) of the according baculovirus P2 stock for 48 hours. Cells were lysed in buffer A by sonication (3x 5 sec on, 20 sec off at 40% amplitude), proteins were purified via Ni-NTA, eluted in buffer B devoid of ATP, aprotinin/pepstatin/leupeptin and PefaBloc but containing 200 mM imidazole. Next, His-tag was removed by overnight TEV cleavage with simultaneous dialysis against buffer A at 4°C. Afterwards, excess His-tagged TEV was removed with Ni-NTA and proteins were further purified over a Superdex 200 10/300 GL column in buffer B.

For expression of Fbxw5, Fbxl2 and Fbxw7, 200 ml of Sf21 cells at 2×10^6^ cells/ml were simultaneously infected with each 1% (v/v) F-box protein- and Skp1-expressing baculovirus P2 stocks for 48 hours followed by lysis in buffer D (20 mM Na-phosphate buffer pH 8.0, 300 mM NaCl, 10 mM imidazole, 1 mM β-mercaptoethanol, 5% glycerol, 1 µg/ml aprotinin/pepstatin/leupeptin and 1 mM Pefabloc) by sonication (3x 5 sec on, 20 sec off at 40% amplitude). Proteins were purified via Ni-NTA, eluted in buffer D devoid of PefaBloc but containing 200 mM imidazole and further purified over a Superdex 200 10/300 GL column in Buffer E (20 mM HEPES pH 7.3, 110 mM K-acetate, 2 mM Mg-acetate, 1 mM EGTA, 1 µg/ml aprotinin/pepstatin/leupeptin).

### Generation of neddylated Cul1 and Cul4A complexes

The bacteria-derived split Cul1A-Cul1B/Rbx1 subcomplex was obtained from B. Schulman. Procedure for generation of neddylated Cul1/Rbx1 and Cul4A/DDB1/Rbx1 from insect cells was modified from Fischer et al., 2011, Li et al., 2005 and Scott et al., 2016^43,72,73^. For Cul1/Rbx1, 200 ml of Sf21 cells at 2×10^6^ cells/ml were infected with 1% (v/v) of the baculovirus P2 stock for 48 hours. Cells were lysed in buffer F (50 mM Tris pH 8, 200 mM NaCl, 5 mM DTT and 30 mM imidazole) by passing the cell suspension twice through an Emulsion Flex-C5 microfluidizer. Proteins were first purified via Ni-NTA, eluted in buffer F containing 200 mM imidazole and further purified over a 1 ml ResourceS cation exchange chromatography column (GE Healthcare) using a 20 ml gradient from 100 mM to 500 mM NaCl in 50 mM 2-(N-morpholino)ethanesulfonic acid (MES) pH 6.5 and 5 mM DTT. Major peak fractions eluting at around 200 mM NaCl were pooled, concentrated and further purified over a Superdex 200 10/300 GL column in buffer G (50 mM Hepes pH 7.4, 200 mM NaCl, 2 mM DTT). His-Cul1/Rbx1 containing fractions were pooled, concentrated and neddylated at 8 µM with 0.1 µM APPBP1/Uba3, 1 µM Ubc12 and 1 mM ATP in neddylation buffer (25 mM Hepes pH 7.4, 200 mM NaCl and 10 mM MgCl_2_). Reaction was started by addition of 20 µM Nedd8, incubated at 30°C for 5 min and stopped by the addition of 10 mM DTT on ice. Components of the neddylation machinery were then removed by another round of Ni-NTA purification. Eluted Nedd8∼His-Cul1/Rbx1 was again concentrated, TEV cleaved overnight at 4°C and further purified over Superdex 200 10/300 GL column in buffer G. Complete neddylation was confirmed via SDS-PAGE (Fig EV2B).

Neddylated Cul4A/DDB1/Rbx1 complexes were generated essentially in the same way, except that the cation exchange step was replace by an anion exchange procedure using a 1 ml MonoQ 5/50 anion exchange chromatography column over a 20 ml gradient ranging from 150 mM to 500 mM NaCl in 50 mM Tris pH 8, 5 mM DTT. Here, Cul4A/DDB1/Rbx1 eluted at around 250 mM NaCl. All following steps including the neddylation reaction and TEV cleavage were conducted as for Cul1.

### Purification of HA-tagged proteins from Hek293T cells

For purification of HA-tagged proteins for *in vitro* ubiquitylation experiments, transiently transfected Hek293T cells from 4x 15 cm dishes were collected 48 hours after transfection, lysed in RIPA buffer (50 mM Tris pH 8.0, 300 Mm NaCl, 1% NP-40, 0.5% Na-deoxycholate, 0.1% SDS, 1 µg/ml aprotinin/pepstatin/leupeptin, 1 mM PefaBloc), cleared by high-speed centrifugation and incubated for 3 hours with 10 µl anti-HA-agarose. Next, beads were washed twice with wash buffer (20 mM HEPES pH 7.3, 110 mM K-acetate, 2 mM Mg-acetate, 1 mM EGTA, 0.1% NP-40, 1 µg/ml aprotinin/pepstatin/leupeptin, 20 mM N-ethylmaleimide (NEM)). Afterwards, beads were washed two more times with wash buffer devoid of NEM. Proteins were eluted by 3 consecutive elutions at 25°C for 10 min with 1 bead volume of elution buffer (20 mM HEPES pH 7.3, 100 mM NaCl, 110 mM K-acetate, 2 mM Mg-acetate, 1 mM EGTA, 0.2 mg/ml ovalbumin, 0.1% NP-40, 1 µg/ml aprotinin/pepstatin/leupeptin, 0.2 mg/ml HA peptide and 50 µM PR619) under constant agitation. Eluates were combined, aliquoted, snap-frozen and stored at −80°C until further use.

### *In vitro* ubiquitylation

*In vitro* ubiquitylation was performed as described previously^24^ with minor alterations. SCF complexes were generated by mixing equimolar amounts of the F-box protein/Skp1 subcomplex with either Cul1/Rbx1 or Cul4A/DDB1/Rbx1 subcomplexes. Ubiquitylation mixes were prepared on ice in a total volume of 20 µl. The reaction was started by adding ATP and stopped by adding 20 µl 2x sample buffer (100 mM Tris pH 6.8, 2% (w/v) SDS, 0.2% (w/v) bromophenol blue, 20% glycerol, 200 mM DTT). For proteins purified from Hek293T cells (Figure 1E), 1-5 µl of HA-tagged candidate substrates were incubated with 170 nM Uba1, 0.5 µM UbcH5b, 0.5 µM Cdc34, 20 µM His_6_-Ubiquitin, 5 mM ATP in the presence or absence of 100 nM SCF^Fbxw5^ in SAB+ buffer (20 mM HEPES pH 7.3, 110 mM K-acetate, 2 mM Mg-acetate, 1 mM EGTA, 0.2 mg/ml ovalbumin, 0.05% Tween-20, 1 µg/ml aprotinin/pepstatin/leupeptin) at 37°C for 2 hours. If not otherwise mentioned, substrates purified from bacteria or insect cells were used at 0.2 µM and in general incubated with 170 nM Uba1, 0.25 µM E2, 75 µM Ubiquitin, 5 mM ATP in the presence or absence of 50 nM SCF^Fbxw5^ in SAB+ buffer. The bacteria-derived split Cul1A-Cul1B/Rbx1 subcomplex obtained from B. Schulman was used for Figure 1, the insect cell-derived Cul1/Rbx1 subcomplex for all other reactions. If not otherwise specified, Sf21-derived MCAK was used as substrate.

### IP and western blot analysis

For co-IP, cells from 1-4 confluent 15-cm dishes were harvested using either a cell lifter or trypsinisation and lysed in 500 µl ice-cold lysis buffer (20 mM HEPES pH 7.3, 110 mM K-acetate, 2 mM Mg-acetate, 1 mM EGTA, 0.2% NP-40 (for Flag-IP) or 0.4% (for Fbxw5-IP), 1 µg/ml aprotinin/pepstatin/leupeptin, 1 mM Pefabloc, 2 mM Na-Orthovanadate, 5 mM NaF). Lysates were cleared by centrifugation 20,000g for 30 min. For Flag-Fbxw5 IPs, 10 µl FLAG-agarose was added for 3 hours under constant agitation. For Fbxw5 IP, 1.5 µg anti-Fbxw5 IgG or unspecific rabbit IgG were added for 1 hour under constant agitation, followed by addition of 10 µl pre-equilibrated Protein A agarose for 2 hours. Beads were washed 4 times with lysis buffer and for Flag-IP, proteins were eluted by 3 consecutive elutions at 25 °C under constant agitation with 3 bead volumes of elution buffer (lysis buffer supplemented with150 mM NaCl and 0.2 mg/ml FLAG-peptide). Eluates were combined and precipitated with 10% trichloracetic acid. Precipitates were washed once with −20°C cold acetone, air-dried and resuspended in 50 µl 1x SDS sample buffer (50 mM Tris pH 6.8, 1% (w/v) SDS, 0.1% (w/v) bromophenol blue, 10% glycerol, 100 mM DTT). For all other IPs, beads were washed 4 times with lysis buffer and proteins were eluted in 40 µl 1x sample buffer.

Since Fbxw5 signals were undistinguishable from background signals upon direct lysis in sample buffer, cells were lysed in 35 µl ice-cold lysis buffer followed by clearance at 20,000 g for 10 min at 4°C. Afterwards, protein concentrations were measured using Bradford assay, adjusted and 30 µl were mixed with 10 µl 4x sample buffer. This procedure was used for all straight western blots shown. MCAK stabilisation at different cell cycle arrests (Fig 4A) upon Fbxw5 knockdown was comparable when obtained by direct lysis in 1x sample buffer (data not shown).

For western blotting, proteins were resolved on 7.5% or 7.5-15% gradient SDS-polyacrylamide gels. After electro-transfer onto PVDF membranes using wet-tank blotting systems (self-made), membranes were stained with coomassie (0.1% coomassie brilliant blue R250 in 50% ethanol and 10% acidic acid) and cut in order to simultaneously detect two proteins from the same blot. After destaining (50% ethanol, 10% acidic acid) and washing with H_2_O, membranes were blocked in PBS-T (140 mM NaCl, 2.7 mM KCl, 10 mM Na_2_HPO_4_, 1.5 mM KH_2_PO_4_, pH 7.5, 0.1% Tween 20) containing 5% skimmed milk powder (Roth) for 2 hours or overnight. Primary antibodies were diluted in PBS-T with 5% milk powder and incubated for 2 hours or overnight. After washing with PBS-T, horseradish peroxidase conjugated secondary antibodies (Jackson Immunoresearch/Invitrogen) were added and proteins were detected with Immobilon Western chemiluminescent HRP substrate and a Fujifilm LAS-4000 Luminescence Image Analyser. In order to detect other proteins on the same blot, blots were incubated over night with the primary antibody of the other protein (different species) together with ∼10 mM NaN_3_ in order to quench signals from the first secondary antibody.

### Pull down experiments

For *in vitro* pull down experiments, purified proteins were mixed in a molar ratio of 5:1 (bait vs. prey) together with competing *E*.*coli* lysate in PD Buffer (20 mM HEPES pH 7.3, 110 mM K-acetate, 2 mM Mg-acetate, 1 mM EGTA, 300 mM NaCl, 20 mM imidazole, 0.1% NP-40, 1 µg/ml aprotinin/pepstatin/leupeptin) and incubated with Ni-NTA agarose for 1h. After 4x washing with PD Buffer, proteins were eluted in 2x sample buffer containing 200 mM imidazole.

### Immunofluorescence

24 hours after siRNA transfection cells were seeded onto 12 mm diameter glass coverslips (Neolab). Cells were fixed in ice-cold methanol for at least 10 min, rehydrated in PBS for 10 min and blocked for 60 min in IF blocking buffer (PBS-T with 2% FBS, 1% BSA, filtered 0.45 µm). Cells were incubated with primary antibodies diluted in IF blocking buffer in a dark and humid chamber for 2 hours. After 3x washing with PBS + 0.1% Tween 20 for 5 min, cells were incubated with Alexa-conjugated secondary antibodies (either anti-mouse or anti-rabbit conjugated either to Alexa488 or Alexa594, respectively) diluted in IF blocking buffer for 1.5 hours in a dark and humid chamber. Hoechst (20 µg/ml) was included with the secondary antibody to stain DNA. After three washes with PBS-T and one wash with PBS, slides were mounted on glass slides (Thermo Scientific) using fluorescence mounting medium. Images (5 stacks 0.5 µm apart) were taken using either a Nikon Ni-E upright microscope equipped with an air objective (Nikon Plan Apo λ 40x NA 0.95) and a DS-Qi2 black and white camera (Fig 7A & Fig EV4B, c) or with a Zeiss Axio Observer fluorescence microscope equipped with oil immersion objectives (LD-Plan-Neofluar 40x/1.3 or Plan-Apochromat 63x/1.4 (Zeiss)), Colibri LED light source, AxioCam MRm camera (all other images).

### Live-cell imaging

RPE-1 cells expressing mNG-MCAK under a doxycycline-inducible promoter were transfected with siRNAs, seeded onto Ibidi 8 Well Glass Bottom µ-slides and induced by adding doxycycline. For pulse-chase experiment, 24 hours post induction and 48 hours post transfection, cells were washed 2x with PBS and incubated in medium without FBS and doxycyclin. Cells were subsequently imaged with a Leica DMi8 Spinning Disk microscope at 40x or 63x magnification (HC PL APO 40x/1.3 Oil or HC PL APO 63x/1.40-0.60 Oil) equipped with a Hamamatsu Orca Flash 4.0 LT. Imaging was performed at 37° C and 5% humidified CO_2_. Telophase cells were chosen for time-lapse imaging and 13 stacks at a distance of 0.5 µm were taken every 20 min. If not enough telophase cells were present, cells in prometa- and metaphase were imaged and quantification started at telophase.

### Image analysis

Image analysis was performed using FIJI^74^. All immunofluorescence images show maximum intensity projections of 5 z-stacks (each 0.5 µm apart) with equal min/max display settings for all images of one experiment. For analysis of MCAK signals at ODF in Fig 4A, total ROIs were generated using a threshold at 200 followed by particle analysis. Total signal intensity of ROIs overlapping with ODF2 signals were measured within maximum intensity projections of the MCAK channel and subtracted by a mean background intensity of a cell-free area multiplied with the area of the corresponding ROI. Analysis of MCAK signals at ODF2 in Fig 5D and Fig EV3B was performed accordingly, except that here ROIs were generated by thresholding after subtracting two maximum intensity projection images subjected to two different Gaussian blurs (σ = 2 or 3 + threshold 25 (Fig 5D) and σ = 3 or 4 + threshold 3 (Fig EV3B)) in order to enable use of same settings for images with highly diverse MCAK signal intensities.

For analysis of time-lapse images, centrosomal mNG-MCAK signals were measured on maximum intensity projections for each time point within a circular ROI (diameter 11 pixels). For background subtraction, extracellular background was measured with the same ROI and the average background intensity for 3 representative movies per experiment was subtracted from the mNG-MCAK centrosomal signal intensity value.

### NanoBRET™ assay

Hela cells were transfected with 2 µg of HaloTag® fusion construct and 20 ng NanoLuc® fusion construct using Fugene® HD (Promega) according to manufacturer’s instructions. Approximately 20 hours later, the cells were harvested, resuspended in NanoBRET™ assay medium (phenol red-free Opti-MEM™ with 4% FCS) and seeded at 2×10^4^ cells/well into white, 96-well flat-bottom tissue culture treated plates (Greiner) in the absence (DMSO vehicle control) or presence of 100 nM HaloTag® NanoBRET™ 618 Ligand. Immediately before measurement, the NanoBRET™ Nano-Glo® substrate (Promega) was added at 10 µM. Luminescence signal was detected at a wavelength range of 415 – 485 nm for the donor and 610 – 700 nm for acceptor signals with the Tecan Spark 10M plate reader. NanoBRET™ ratio (BU) was calculated by dividing the acceptor signal by the donor signal. The corrected NanoBRET™ ratio was calculated by subtracting the DMSO vehicle control sample from the experimental sample with cells treated with HaloTag® NanoBRET™ 618 Ligand.

### Incucyte

Cells were seeded on 12-well plates 24 hours before imaging with an Incucyte® S3 Live-Cell Analysis System. Images were taken using the 20x air objective. H2B-mRuby signals were used to count all cells automatically using the same settings for all images via the Incycyte software. Mitotic cells were counted manually and the percentage was afterwards calculated.

### Plasmid generation

All plasmids used in this study are shown in Appendix Table 1 and were generated by standard cloning techniques either via site-directed mutagenesis, Gibson assembly^75^ or restriction enzyme-based strategies (New England Biolabs) using *E*.*coli* strain DH5α. Oligonucleotides for PCR amplification were obtained from Sigma Aldrich. For each construct the relevant open reading frame was completely verified by Sanger sequencing (GATC/Eurofins Scientific). Plasmid name indicates vector backbone followed by gene of interest with or without tag or mutations. All genes are of human origin, except for plasmids pBigBac-HisCul4A/His-DDB1/His-mmRbx1 and pBigBac-HisCul1/His-mmRbx1, in which Rbx1 is derived from *Mus musculus*. pFastBac1-Skp1ΔΔ contains human Skp1 with deletion of 2 loops (aa 38-43 and 71-82)^76^. pBigBac-HisCul4A/His-DDB1/His-mmRbx1 contains Cul4A (aa 38-759), DDB1 (aa 1-1140) and mmRbx1 (aa 12-108)^72^. pBigBac-HisCul1/His-mmRbx1 contains full-length Cul1 and mmRbx1 (12-108). pcDNA3.1-HA-CSPP1 contains CSPP1 (878-1256). All constructs generated in this work contain the corresponding canonical sequence (according to http://www.uniprot.org), except for Kif2A constructs (isoform 1, identifier: O00139-1), ODF2 constructs (isoform 3, identifier: Q5BJF6-3).

## Expanded View Figure legends

**Dataset EV1**

Table contains data of protoarray screen. First spreadsheet shows normalised and adjusted intensity values of all candidates, second spreadsheet values of all proteins spotted on the protoarray. The third spreadsheet shows raw data obtained within the protoarray screen and the fourth spreadsheet shows the GO analysis (cellular components) using the DAVID webtool.

**Expanded View Figure 1 (related to Figure 1)**

**A**. Scatter plots comparing intensities of duplicate SCF^Fbxw5^ (left) and control (right) samples. Dotted lines indicate the cut-off for candidate substrates (500 AU). **B**. Comparison of all protoarray signal intensities probed with or without SCF^Fbxw5^ complexes. **C**. Co-IP of endogenous HGS, Sec23B, CSRP2 and GSTP1 with exogenously expressed Flag-Fbxw5 as in C. **D**. Co-IP of endogenous HGS and Sec23B with endogenous Fbxw5 as in C, except that anti-Fbxw5 antibodies were used instead of anti-Flag. Source data are presented in Dataset EV1.

**Expanded View Figure 2 (related to Figure 3)**

**A**. Coomassie-stained SDS-PAGE of neddylation enzymes purified from *E*.*coli* (left) and Cul1/Rbx1 and Cul4A/DDB1/Rbx1 complexes purified from Sf21 cells (right) together with Ube2G1 from *E*.*coli*. **B**. Coomassie-stained SDS-PAGE of Cul1 (left) and Cul4A (right) complexes before and after neddylation (N8 = Nedd8). **C**. *In vitro* ubiquitylation. 0.2 µM MCAK was incubated with 75 µM ubiquitin (untagged or His_6_-tagged), 170 nM E1, 1 µM of UbcH5b, 0.15 µM Fbxw5/Skp1 and 0.15 µM Cul1∼Nedd8/Rbx1 at 30°C for 45 min followed by western blotting using anti-MCAK (top) or anti-Fbxw5 antibodies (bottom) for detection. **D**. Experiment as in C, except that 0.2 µM His_6_-Eps8 was used and incubation time was 90 min at 30°C. **E**. Experiment as in Fig 3B with Cdc34 as E2, except that untagged, His_6_-tagged or Flag-tagged ubiquitin was used. **F**. Time course experiment as in Fig 3C, except that 0.25 µM UbcH5b was used and reaction was carried out for 60 min at 30°C. **G**. 0.2 µM MCAK was incubated with 170 nM E1, 1 µM UbcH5c, 1 µM Ube2G1 and 0.2 µM Fbxw5 premixed either with 0.2 µM Cul1∼Nedd8/Rbx1 (left) or 0.2 µM Cul4A∼Nedd8/DDB1/Rbx1 (right) at 30°C for 15 min. **H**. Coomassie-stained SDS-PAGE of F-box proteins, each purified from Sf21 cells in complex with Skp1 using same procedures. **I**. Experiment as in Fig 3E, except that 0.25 µM of UbcH5b was used and reaction was carried out at 30°C for 60 min. **J**. comparison of MCAK ubiquitylation purified from *E*.*coli* or Sf21 cells. Experiment as in Fig 3B, except that for blots 1&2 1 µM of UbcH5b and 0.15 µM SCF^Fbxw5^ was used. For blot 3, 0.25 µM Cdc34 and 0.05 µM SCF^Fbxw5^ was used. **K**. Coomassie-stained SDS-PAGE of different ubiquitin variants. Methylated ubiquitin (last lane) was obtained from BostonBiochem, all others were purified from *E*.*coli*. **l**, Experiment as in Fig 3G with Cdc34 as E2, except that Kif2a purified from Sf21 cells was used instead of MCAK and reaction was carried out for only 5 min at 30°C.

**Expanded View Figure 3 (related to Figure 5)**

**A**. Comparison of mNG-MCAK localization with endogenous MCAK by IF. Top panels show mNG-MCAK signals of methanol-fixed and Hoechst-stained cells in different cell cycle stages. Bottom panel shows endogenous MCAK signals obtained by IF with anti-MCAK antibodies in methanol-fixed and Hoechst-stained wild type RPE-1 cells. Scale bar = 5 µm. Note that different min/max settings have been applied for better visualisation **B**. RPE-1 cells were treated with the indicated siRNAs for 48 hours and arrested with 75 ng/ml nocodazole for 5 hours. Mitotic cells were shaken off, washed twice with PBS, plated on coverslips in serum-free medium and fixed at the indicated time points with methanol followed by immunofluorescence using the indicated antibodies. For zero hour time point, cells were directly seeded on coverslips after siRNA treatment and harvested after 5 hours nocodazole-arrest. Left: Maximum intensity projections of representative images. Arrows indicate ODF2 signal. Scale bar = 5 µm. Note that for 0 hour time point different min/max display settings were applied as for the other time points for better visualisation (settings are always kept the same within each time point and no adjustment was applied for quantification). Right: Quantification. For time point 0h, all ROIs (obtained by thresholding) were considered (kinetochore signals). For time points 4, 18 and 24 hours only signals co-localising with the ODF2 spot were considered (centrosomal signals). Error bars show standard error of the mean of 3 independent experiments covering in total more than 150 cells. Asterisks indicate p-value of a Mann Whitney test comparing Fbxw5 knockdown with control samples for each time point. **C**. Polyclonal RPE-1 cells were treated with the indicated siRNA for 24 hours, split on Ibidi 8 Well Glass Bottom µ-slides and mNG-MCAK expression was induced with 10 ng/ml doxycyclin for another 24 hours. Cells in prometaphase were imaged using spinning disk microscopy. Left: Representative images. ROI used for measuring signal intensities is indicated. Scale bar = 5 µm. Right: Quantification of total mNG-MCAK signal intensity in prometaphase cells of 3 independent experiments each covering more than 30 cells. Error bars indicate standard error of the mean, asterisks p-value of a Student’s t-test. Source data for B are presented in Source Data Table 2, for C in Source Data Table 3.

**Expanded View Figure 4 (related to Figure 7)**

**A**. Polyclonal RPE-1 cells stably expressing H2B-mRuby2 under a CMV promoter were transfected with the indicated siRNA for 48 hours and then imaged using Incucyte® S3. Chart shows the percentage of mitotic cells of 3 experiments, each covering more than 2500 cells. **B**. Quantification of the length of remaining cilia of RPE-1 cells upon mNG-MCAK overexpression (same samples as in Fig 7A). Long black line shows mean intensity, error bars show standard error of the mean of 3 independent experiments each covering more than 40 cilia and asterisks indicate p-value of a Student’s t-test comparing the indicated samples. **C**. Correlation between MCAK signal intensity and cilia length of same samples as in B. MCAK signal intensity was measured within a circular ROI at basal bodies with a diameter of 1.65 µm and plotted against the according length of the remaining cilia. Solid line shows simple linear regression and dashed line the 95% confidence bands. R^2^ and the p-value of a Pearson correlation analysis are shown. **D**. Quantification of the length of remaining cilia of RPE-1 cells upon siRNA treatment (same samples as in Fig 7B). Long black line shows mean intensity, error bars show standard error of the mean of 4 independent experiments each covering more than 30 cilia and asterisks indicate p-value of a Student’s t-test comparing the indicated samples. **E**. Polyclonal RPE-1 cells stably expressing H2B-mRuby2 under a CMV and mNG-MCAK under a doxycycline-inducible promoter were treated with 20 ng/ml doxycycline for 24 hours in full serum and then imaged using the Incucyte S3. Chart shows the percentage of mitotic cells of 3 experiments, each covering more than 3500 cells. Source data are presented in Source Data Table 3.

